# Epigenetic regulation affects fertility in *Drosophila*: toward the production of infertile models

**DOI:** 10.1101/317123

**Authors:** Hidetsugu Kohzaki, Maki Asano, Yota Murakami, Alexander Mazo

## Abstract

We have revealed that the chorion gene clusters amplify by repeatedly initiating DNA replication from chorion gene amplification origins in the response to developmental signals, through the transcription factors in *Drosophila* ovarian follicle cells. Orc1, Orc2, and Cdc6 are forms of DNA replication machinery, which are conserved from yeast to humans; and Orc1 and Orc2 mutants are lethal. Overexpression of Orc1 or Orc2 (subunits of the origin recognition complex) led to female sterility, but overexpression of Cdc6 (an Orc family member) or GFP did not. We propose that DNA replication machinery contributes to development.

Recently, we found that H3K4 was trimethylated at chorion gene amplification origins, but not at the Act1 locus. Overexpression of Lsd1H3K4 dimethylase and Lid H3K4 trimethylase are female sterile but not a Lid mutant. These results showed that epigenetic regulation affected fertility. Screening strategies using *Drosophila* flies could also lead to the development of drugs that reduce sterility and epigenetic effects related histone modification.

**Summary statement:** There are approximately 470,000 infertile individuals in Japan. We knockowned the prereplicative complex components and demethlases during *Drosophila* ovary development. In these drospohila, we could be the model of infertile.

## Introduction

In *Drosphila*, different DNA replication systems were used during their development process. In *Drosophila* oogenesis, not only does the endoreplication occur in the nurse cells and follicle (fc) cells, but chorion gene amplification also take place in fc cells (Lin et al., 1994; Khoury Christianson, A. M., King, D. L., Hatzivassiliou, E., Casas, J. E., Hallenbeck, P. Calvi et al., 1998; Royzman and Orr-Weaver, 1998; Lee and Orr-Weaver, 2003; Claycomb et al., 2004; Spradling and Orr-weaver refs). Amplification of *Drosophila* chorion genes is necessary for eggshell formation, and mutations that disrupt amplification, such as Cyclin E, Orc2 (Landis et al., 1997; Loupart et al., 2000), chiffon (Dbf4) Landis and Tower, 1999), humpty dumpty, Mcm6 (Schwed et al., 2002) and so on, cause female sterility. Orc2, Orc5 and Orc6 (Loupart et al., 2000; Pflumm and Botchan, 2001; Balasov, et al., 2009) showed strong S-phase defects. Surprisingly, dE2F1, dDP (Royman et al., 1999), dE2F2 (Cayirlioglu et al., 2001; Cayirlioglu et al., 2003), Rbf (Basco et al., 2001) and Myb complex (Beall et al., 2002; Beall et al., 2004) are necessary for this process (Cayirlioglu et al., 2003). We proposed that the chorion gene cluster amplify by repeatedly initiating DNA replication from chorion gene amplification origins (Lu et al., 2001; Zhang and Tower, 2004) in the response to developmental signals through the transcription factors in ovarian fc cells (kohzaki & Murakami 2007).

Orc1 is a large subunit of Orc (<u>O</u>rigin <u>r</u>ecognition <u>c</u>omplex) complex and functions as a main menber of pre-RC. In *Drosophila*, Orc1 level are transcriptionally upregulated by E2F (Asano and Wharton, 1999) and then down regulated by APC (<u>a</u>napahse <u>p</u>romoting <u>c</u>omplex) for proteolysis (Araki et al., 2005). Not only Orc1 but also Orc2 in the salivary and ovian fc cells are not requierd for endoreplication (Park and Asano, 2008). When Orc1 was absent, most amplification diminished, whereas when Orc1 was overexpressd to fill the entire nuclei, DNA replication occurred throughoyt entire nuclei. These results suggested that Orc1 is a limiting factor at least in some tissues. On the other hand, Orr-Weaver et al. showed that amplification of at one locus does not require Orc for intiation demonstrated that Orc-independent DNA replication is used multiple modes of replication in intact organisim (Kim and Orr-Weaver, 2011).

We’ve reserached where the initiation of DNA replication start and what at first makes it to begein. In this study, we uncoverd the regulation of signal transduction and DNA reolication, especially, about EcR and its cofactor TRR H3K4 trimethyrase at the step of Orc1 loading.

## Results

### Ecdyson contols Orc1 loading at gene amplification locus neccessory for *Drosophila melanogaster* oogenesis

Steroid hormons have important functions in animal development. In *Drosophila*, ecdysone triggers moulting, metamorphosis and oogenesis through its effect on gene expression network (Bender et al., 1997; White et al., 1997; Buszczak et al., 1999; Tzolovsky et al., 1999; Dej and Spradling 1999; White et al., 1999; Carney and Bender, 2000; Arbeitman et al., 2002; Li and White, 2003; Davis et al., 2005, Kohzaki et al., 2018a). Ecdysone works by binding to a nuclear receptor, EcR (Koelle et al., 1991). The EcR heterodimerizes with the retinoid X receptor ortholog Ultraspiracle (USP), which acts as a general heterodomer partner for the class of factors repsentated by EcR (Hall and Thummel, 1998; Arbeitman and Hogness, 2000; Ghbeish et al., 2001; Hu et al., 2003). Both partners are required for binding to ligand or their target DNA. This heterodimer activate EcR response gene expression by recruiting co-regulators. Mazo et al. showed TRR, which is a histone methyltransferase capable of trimethylating lysine 4 of histone H3 (H3-K4), function as a coactivator of EcR by altering the chromatin structure at ecdysone-responsive promoter (Sedko et al., 1999; Sedkov et al., 2003). Also, Gerbi et al. showed that Ecdysone induced gene amplification in *Sciara coprophila* DNA puff II/9A (Liang et al., 1993; Foulk et al., 2006).

Recently we could find Orc1binds to ACE and Ori *β* directly (Kohzaki and Asano, 2016) using flies having a single copy *orc1* promoter *orc1*+-GFP-9myc transgene (Orc1-GFP9myc) (Park and Asano, 2008). In this Fly, Orc1-GFP expressed endoginous Orc1 similally (Fig.1A and C) and localized follicle cells (Fig.1B and D). We’ve searched whether responsing elements are or not around gene amplification locus. We can detect several EcR putative binding sites (Fig.2A). in these flies ovaries, we performed ChIP assay using EcR-C monoclonal antibody and TRR polyclonal antibody (Sedkov et al., 2003). We detected ACE3, ori-*β* and ACE1 signal. This data showed that EcR and TRR localizes on ACE3 and ACE1 (Fig.2). The amounts of ACE1, ACE3, ori-β PCR products obtained using EcR, TRR, Tri-Me and Myc (for Orc1) antibody relative to Di-Me antibody were significant, respectively (T test, *p*>0.05) (Kohzaki and Murakami, 2007). Also, the amounts of ACE1, ACE3, ori-*β* PCR products obtained using EcR, TRR, Tri-Me and Myc (for Orc1) antibody were no statistical difference, respectively (T test, *p*>0.05). These data showed that EcR, TRR and Orc1 could form complex in a situation such as the initiation of DNA replication.

**Figure 1.**
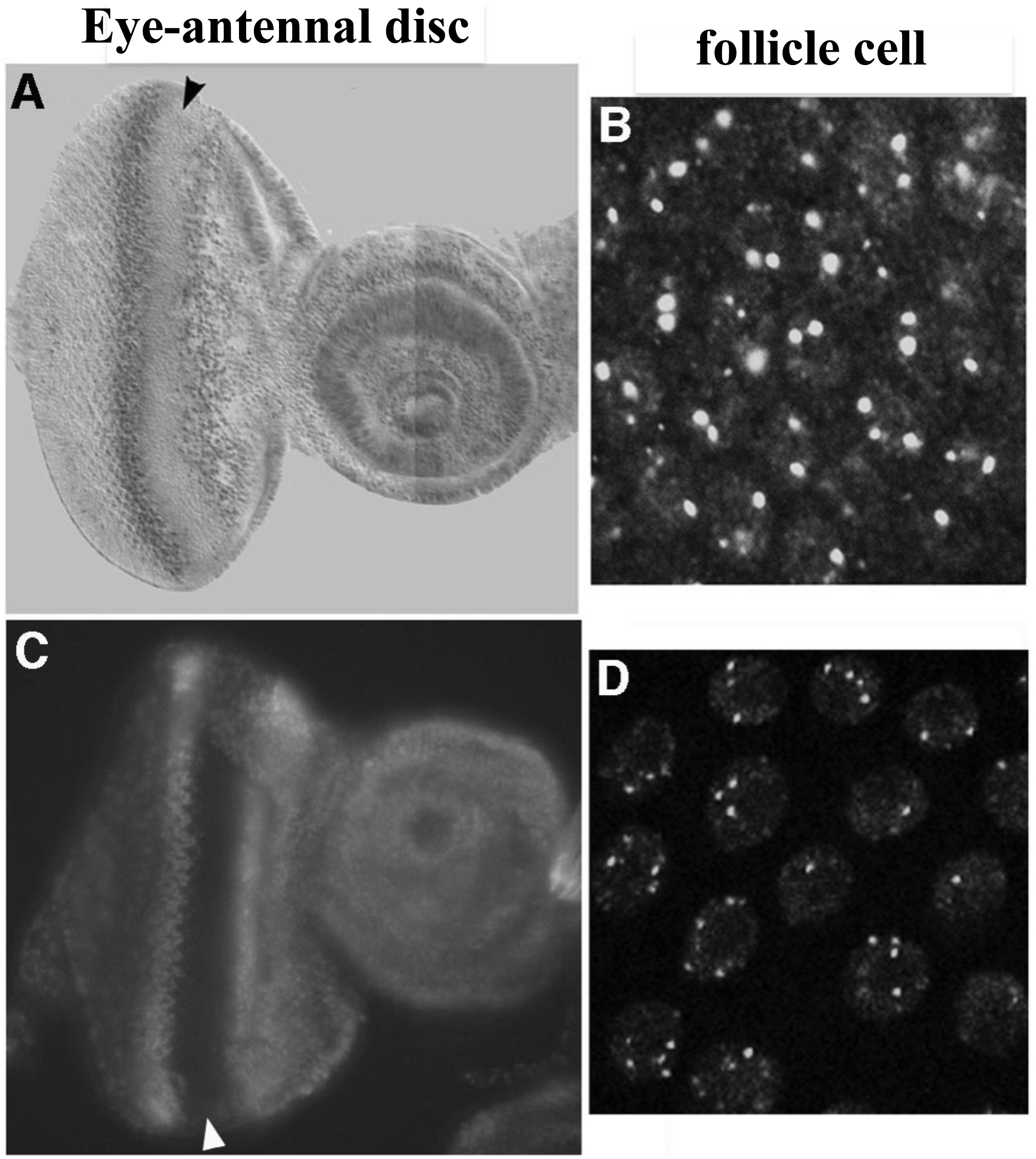
Distribution of Orc1 and Orc1-3HA-GFP. (A, B) Antibody straining reveals the distribution of endogenous Orc1 using anti-Orc1. (C, D) Fluorescent signals from Orc1-GFP9myc driven by *orc1* promoter.

**Figure 2.**
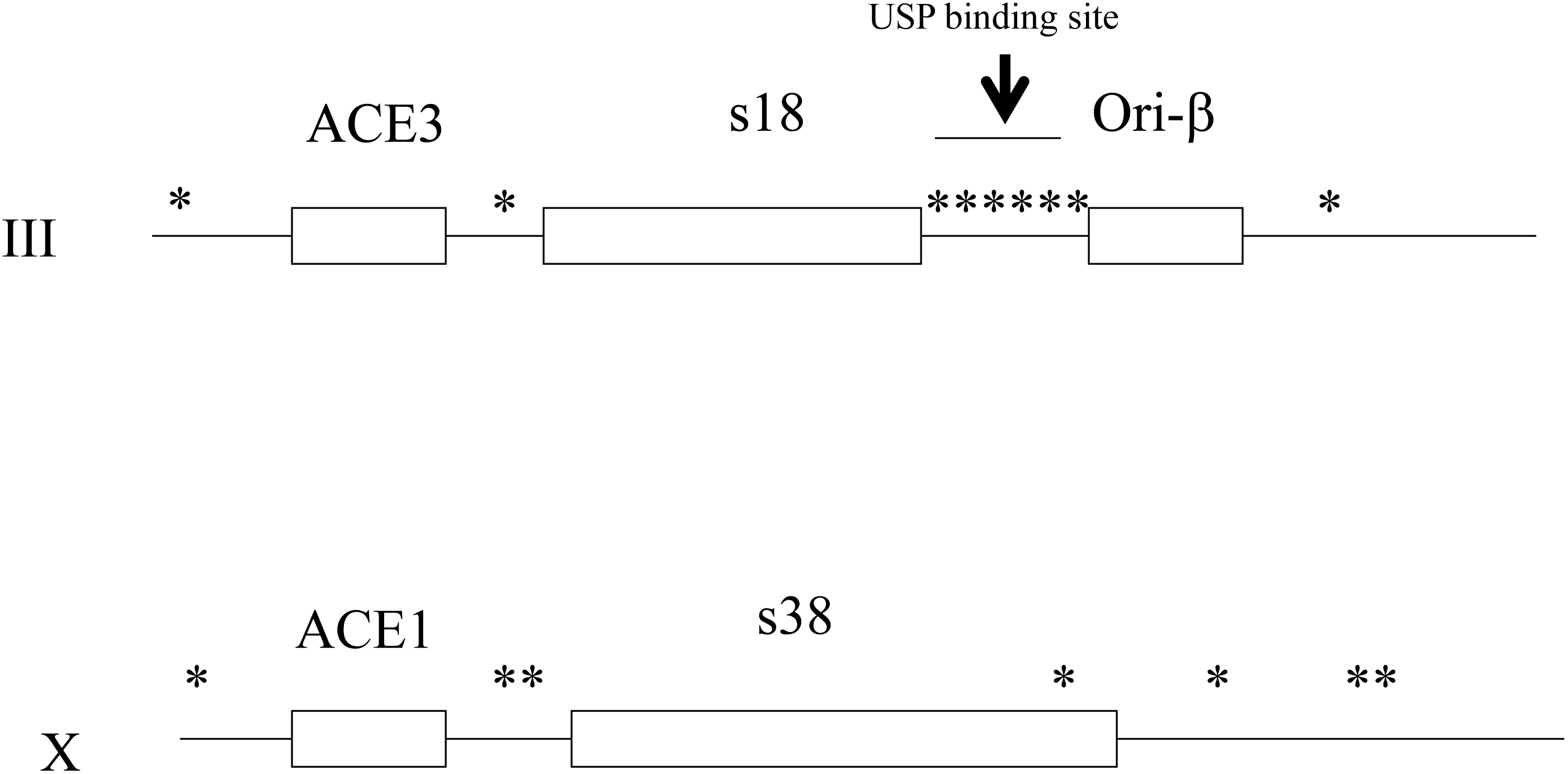
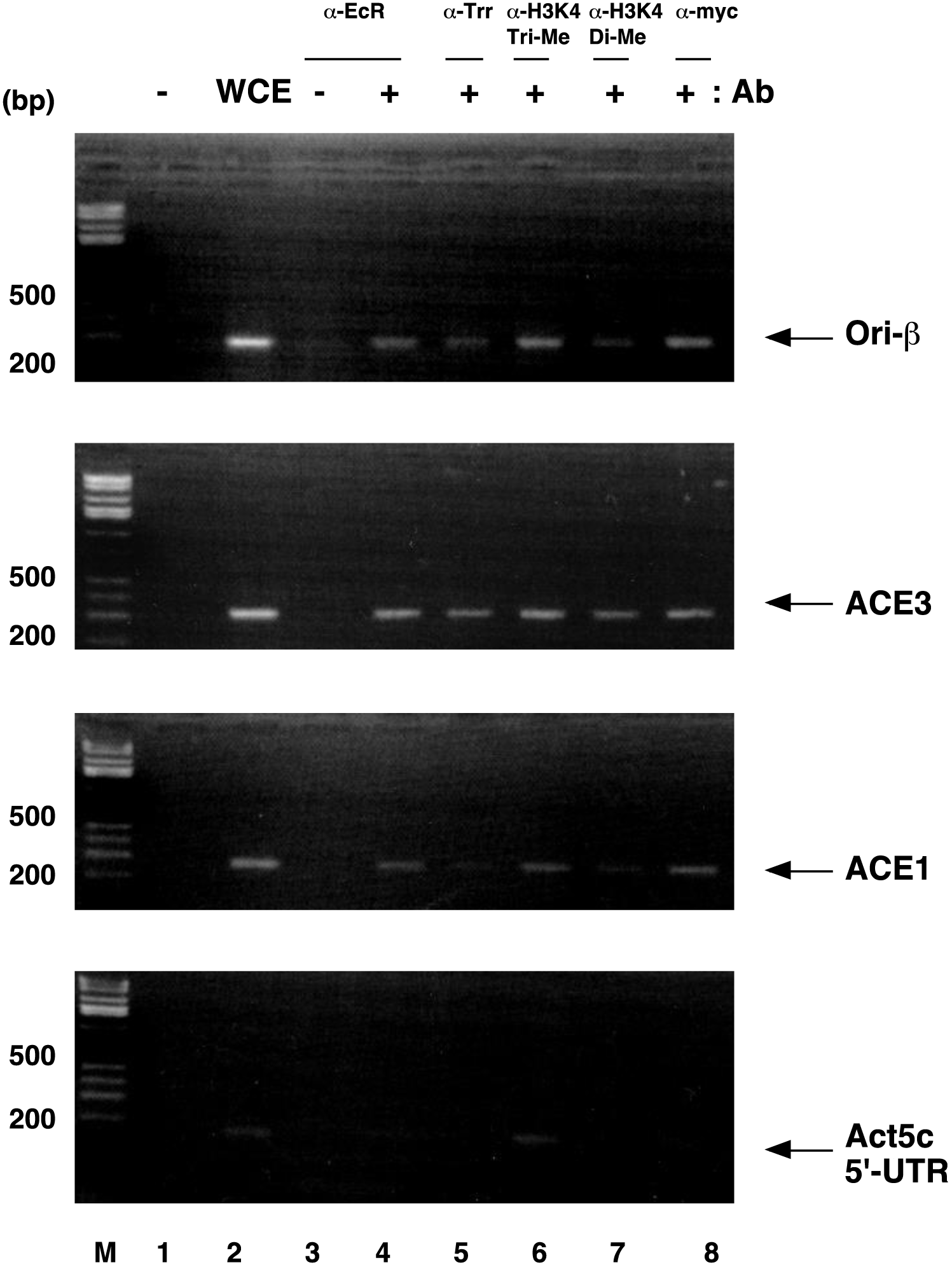
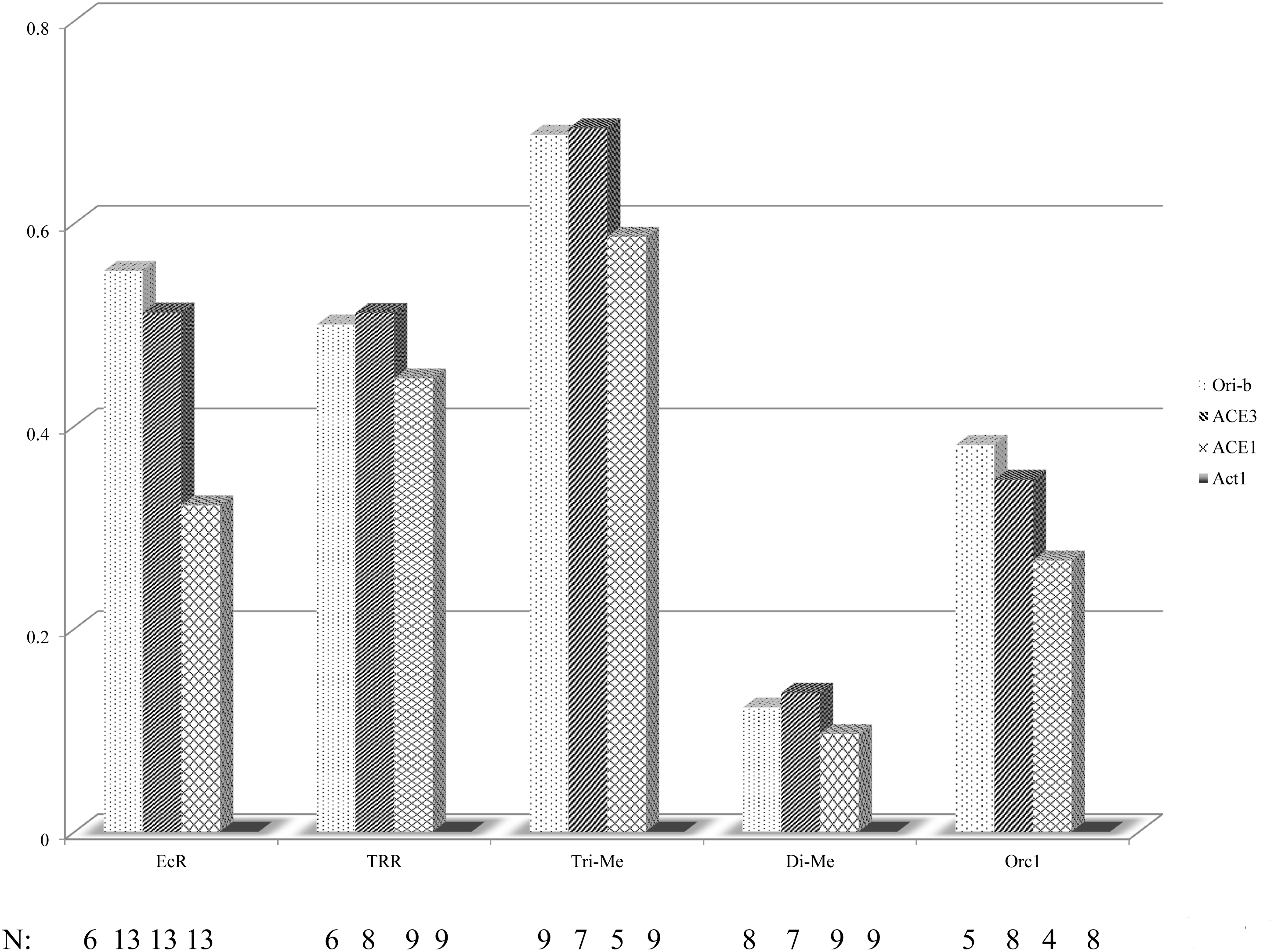
(A) Schematic representation of EcR binding site. Asterisks were putative binding sites. USP binding site was reported previously (Shea et al., 1990; Khoury Christianson et al., 1992; Mariani et al., 1996). (B) Association of EcR, TRR, Trimethylatrd Histone H3K4, dimethylated Hitone H3K4 and Orc1-GFPmyc with chorion gene elements in vivo. ChIP assay were performed as described in Methods with anti-EcR, anti-TRR, anti-Trimethylatrd Histone H3K4, anti-dimethylated Hitone H3K4, and anti-myc antibody (lane 4-8). DNA was amplified using PCR primers specific ACE1, ACE3, Ori-*β* and Act5C 5’-UTRas described previously(Austin et al., 1999; Kohzaki and Asano, 2016) 1A. these primers were also used to amplified DNA isolated from whole cell extracts before immunoprecipitation (WCE). The experiments were performed to confirm reproducibility as describe in Fig.2B. M, DNA size markers; lanes 1 and 2, PCR controls with no DNA (lane 1) or with Whole cell extract (lane 2). (C) quantitation of ChIP assay. The experiments were performed several times(N). The amounts of PCR products obtained from WCE are 1.0 (lane 2 in Fig.2B).

On the other hands, we planned to eliminate ecdyson signal. Four Ecdyson recepter isoform are isolated (Cherbas et al., 2003; Davis et al., 2005). Each isoform has function at each tissue. We overexpressed each ecdyson recepter at the follicle cells. As shown in Fig.3, overexpression of ecdyson recepters except EcR.B1 leaded to female sterile but not EcR mutant F645A which did not have the transcription activity. These data showed that ecdyson may regulate the gene amplification directly through the the transactivation.

**Figure 3.**
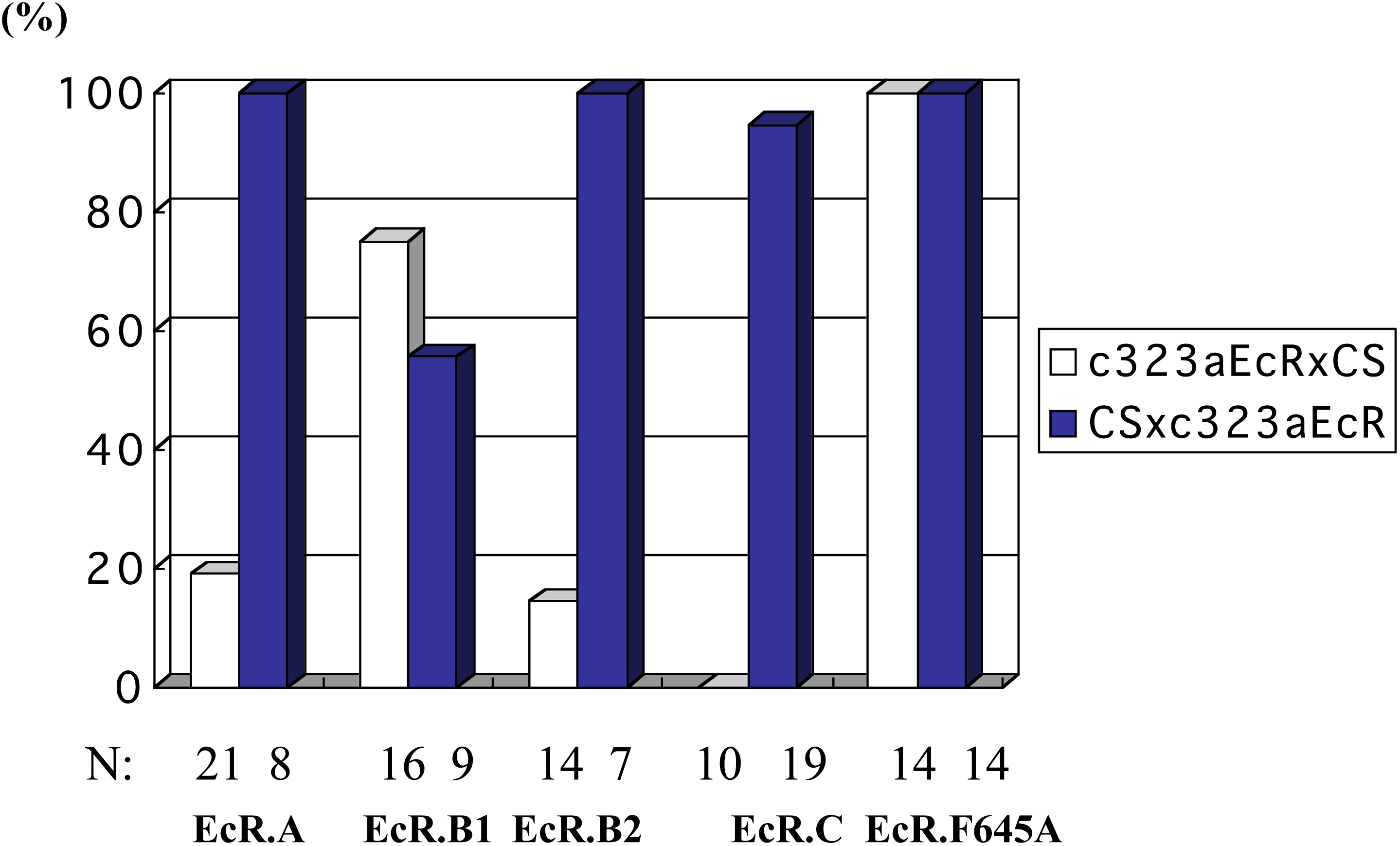
overexpression of dominant negative forms of EcR led to femal sterile. Females of *c323a-Gal4* driver was crossed with male flis harboring UAS-EcR isoform transgene (a responder). The progenies (males or females) having the indicated responder and *c323a* were crossed to wildt ype fly Carton S. All responders were tested: if no bar is visible, there were no progeny with wild type phenotype including growth rates and rate of larvae emergency. The experiments were performed several times(N). The numbers of born flies from CS (male) x *c323a* expressing EcR families (female) are 100% (blue bars in Fig.3).

**Figure 4.**
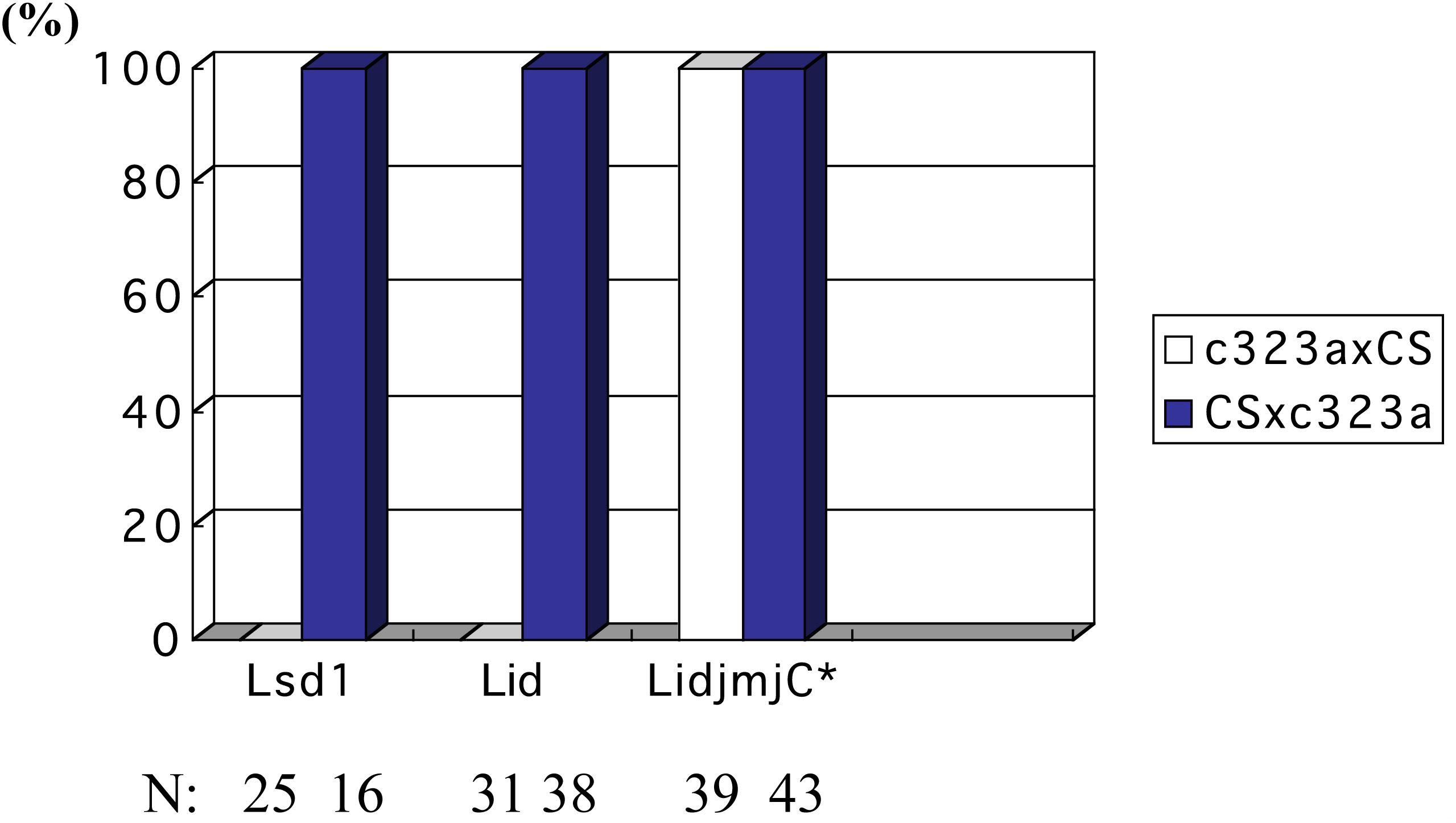
overexpression of Lsd1 and Lid led to femal sterile but not LidjmjC*. Assays were performed as described in Figure 3 with flies harboring UAS-Lsd1, UAS-Lid and UAS lid jmjC* transgenes. The experiments were performed several times (N). The numbers of born flies from CS (male) x *c323a* expressing Lsd, Lid or LidjmjC* families (female) are 100% (blue bars in Fig.3).

Next question is whethere Histon H3K4 is converted to trimethylated form (Tri-Me H3K4) by TRR. TRR was identified as a Set domain protein in *Drosophila* and is highly homologous to drosophila TRITHORAX protein amd to its human homologue, ALL-1/HRX. TRR mutant, trr1 and trr3 was isolated and showed the embryonic lethal (Sedko et al., 1999). Trimethylated H3K4 is associated with transcriptionally active genes in eukaryotes (Metzger and Schüle, 2007; Sims and Reinberg, 2007). We checked the localization of Tri-Me H3K4 around gene amplification locus. Actin 5C is transcribed during early embryogenesis. In fact, Actin 5C 5’UTR was trimethlated, but not dimethyated. We found the localization of Tri-Me H3K4 around ACE3, ori-*β* and ACE1. H3K4 was dimethylated, too (Fig.2). These data showed that gene amplification locus is euchromatin structure. The gene amplification locus codes the genes for choriogenesis. The gene amplification might be induced by ecdyson signal (Fig.2). We observed Tri-Me H3K4 of gene amplification and TRR loading in fc cell. For we investigate biological significance of Tri-Me H3K4, we tried to overexpress H3K4 demethylase Lsd1 or Lid in fc cells. Many groups isolated H3K4 demethlases and trimethylases competitively among same spoeces (Shi et al., 2004; Shi et al., 2005; Christensen et al., 2007; Lan et al, 2007; Liang et al., 2007; Seward et al., 2007; Stefano et al., 2007; Iwase et al., 2007; Tahiliani et al., 2007). In *Drosophila*, Lsd1 is H3K4 dimethylase (Stefano et al., 2007) and Lid is H3K4 trimethylase (Eissenberg et al., 2007; Lee et al., 2007; Secombe et al., 2007). As shown in Fig.3, Lsd1 or Lid overexpressed female is sterile. In the other hand, female expressing Lid jmjC* which has a mutation on active site and Cdc6, GFP (Kohzaki et al., 2018a) are normal. These data showed that H3K4 Tri-Me is essential for gene amplification.

Orc1 is a large subunit of Orc complex, which is the initiator of chromosomal DNA replication. Orc1 is the key player because Orc1 binds to chromatin by BAH domain and is degraded in cell cycle dependent manner (Asano et al., 1996; Asano and Wharton, 1999; Kohzaki and Asano, 2016). And for functional Orc complex formation, Orc core complex, Orc2-5, is necessary for Orc1 (Austin, R. J., 1999; Chesnokov et al., 1999; Pflumm and Botchan, 2001; DePamphilis et al., 2006). In Fig. 2, Orc1 is loaded on gene amplification origin, too. These data showed that developmental signals could regulate gene amplification. We supeculate this reaction is coupled to transcription. We considered whether EcR-TRR signal could regulate the Orc1 loading directly. We planned the overexpression of Orc1 in fc cells using fc specific Gal4 driver (Fig. 5), *c323a* for bypass of developmental signal. These flies were female sterile (Kohzaki et al., 2018a).

**Figure 5.**
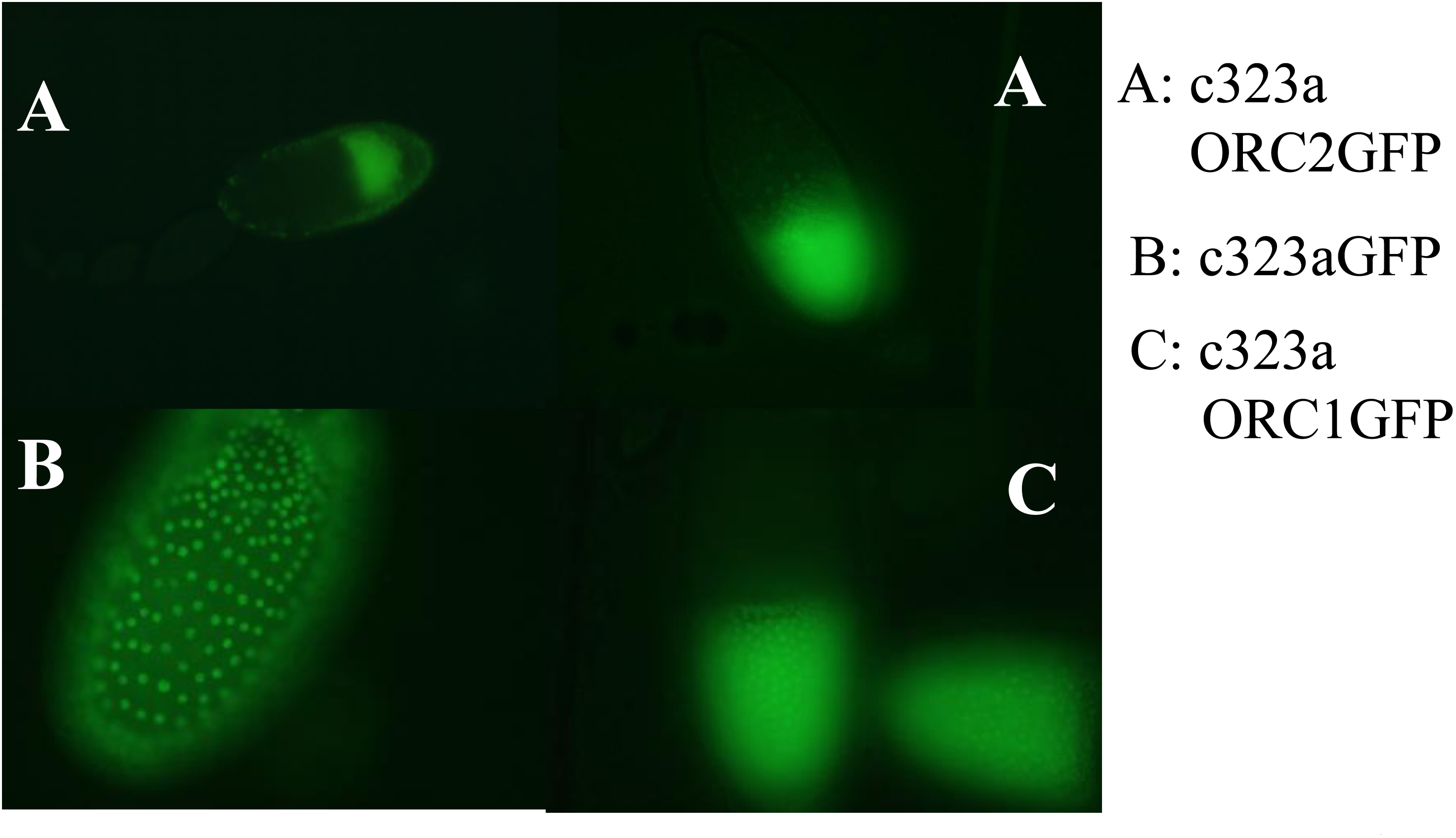
overexpression of Orc1 and Orc2 in fc cells. They led to femal sterile but not GFP.

In summary, these data suggest that Ecdyson signal could determine which DNA replication origin to fire (Fig. 6).

**Figure 6.**
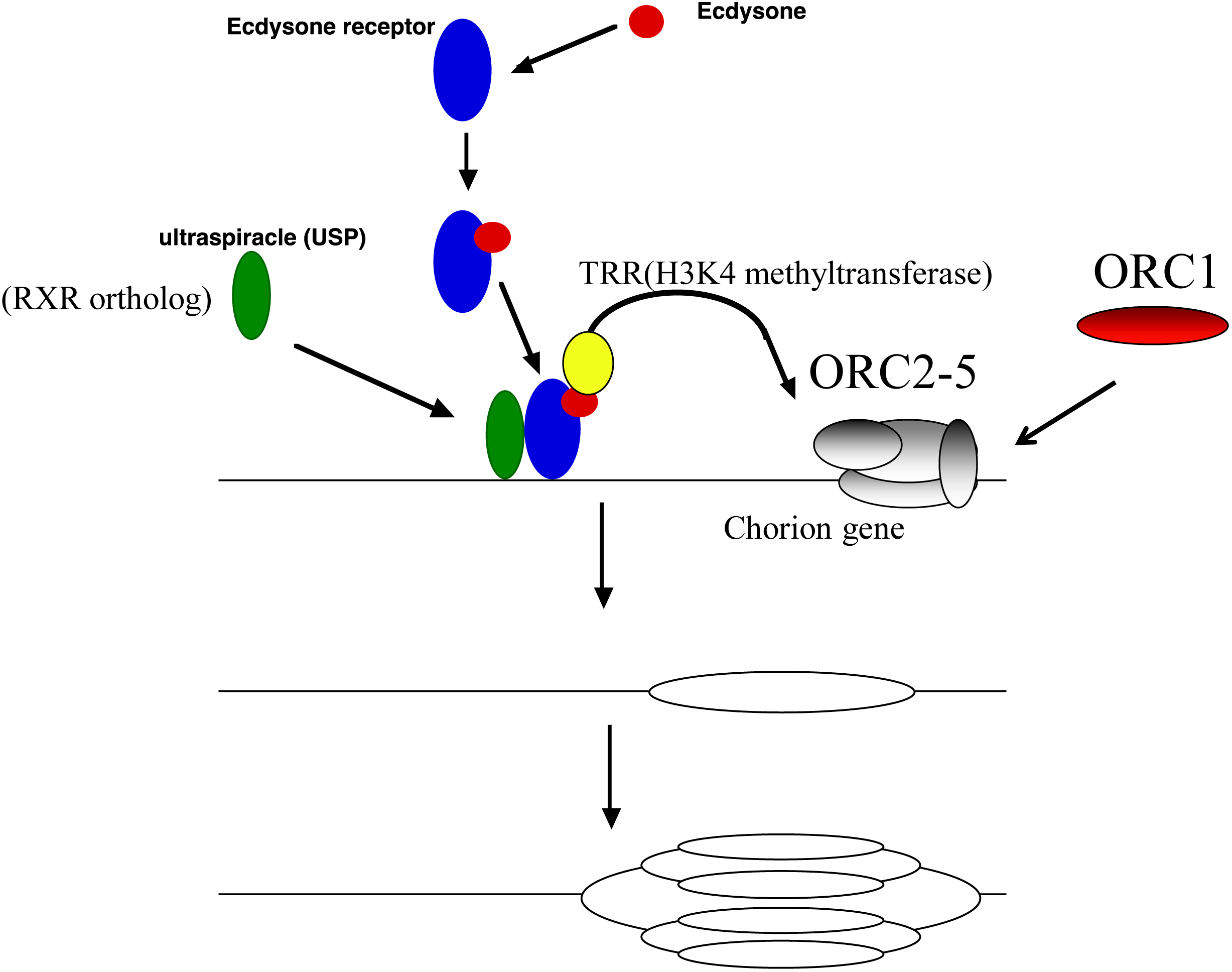
Schematic representation of the putative machanism in which Ecdysone signals could regulate the chorion gene amplification.

## Discussion

In this issue, we showed that EcR regulated chorion gene amplification through the activity of H3K4 trimethylase, TRR. Conversely, overexpression of H3K4 demethylase, Lsd1 and Lid showed the female sterile but not the jmjC mutant form, LidjmjC*. Because EcR mutant (Buszczak et al., 1999; Dej and Spradling, 1999; Carney and Bender, 2000), TRR mutant (Sedkov et al., 1999) and Orc2 mutant (Landies et al., 1997) are growth defect before the chorion gene amplification occurs, EcR signals could direct the gene amplification. The heterodimer partner USP was identified originally as chorion factor 1, which bound to chorion s15 cis-regulatory element (Shea et al., 1990; Khoury Christianson et al., 1992; Mariani et al., 1996). This region includes ori-*β* and our putative EcR binding sites (Fig. 2A). We proposed that EcR-USP-TRR bound to the region between S18 and ori-*β*.

We showed that transcription factors regulates Orc loading and initiation of DNA replication by chromatin modification. in *S. cerevisiae* (Kohzaki, et al., 1999) and *Drosophila* (Kohzaki and Asano, 2016). It was shown that the initiation of gene amplification was associated with histone H3 and H4 hyperacetylation and H1 phosphorylation in *Drosophila* (Hartl et al., 2007). Indeeed. in *Drosophila* fc cells, tetering Rpd3 or Polycob proteins to the origins decreasd its activity, whereas tethering the Hat1 homologue, Chameau acetyltransferase increased origin activity (Aggarwal and Calvi, 2004). This assay used artificial tecnique, tethering. It remains unclear in vivo. Orr-Weaver and her collegues showed that dE2F-dDP-Rbf interact with DmOrc and dE2F1 and DmOrc were bound to chorion gene amplification locus in vivo (Basco et al., 2001). As they did not identify the mutation disrupted the interaction and where is the E2F binding site, it remain uncovered that the they bound directly or indirectly.

Recently, Overexpressed Orc1 is localized in fc cell nuclei. But These flies are female sterile. GFP having NLS and Cdc6 expressing flies are no phenotype. Orc2 expressing flies are female sterile, too. But this phenotype is mider than Orc1overexpressing fies (Kohzaki et al., 2018a). Overexpression of Orc2 led to titrate out the other DNA replication machinary becouse Orc2 is a core member, Orc2-5 (Chesnokov, et al., 1999) (Fig.7A). On the other hand, Orc1 is maybe a limiting factor. These result indicated that Orc1 can bypass developmental signal and dysregulate origin fireing (Fig.7B).

**Figure 7.**
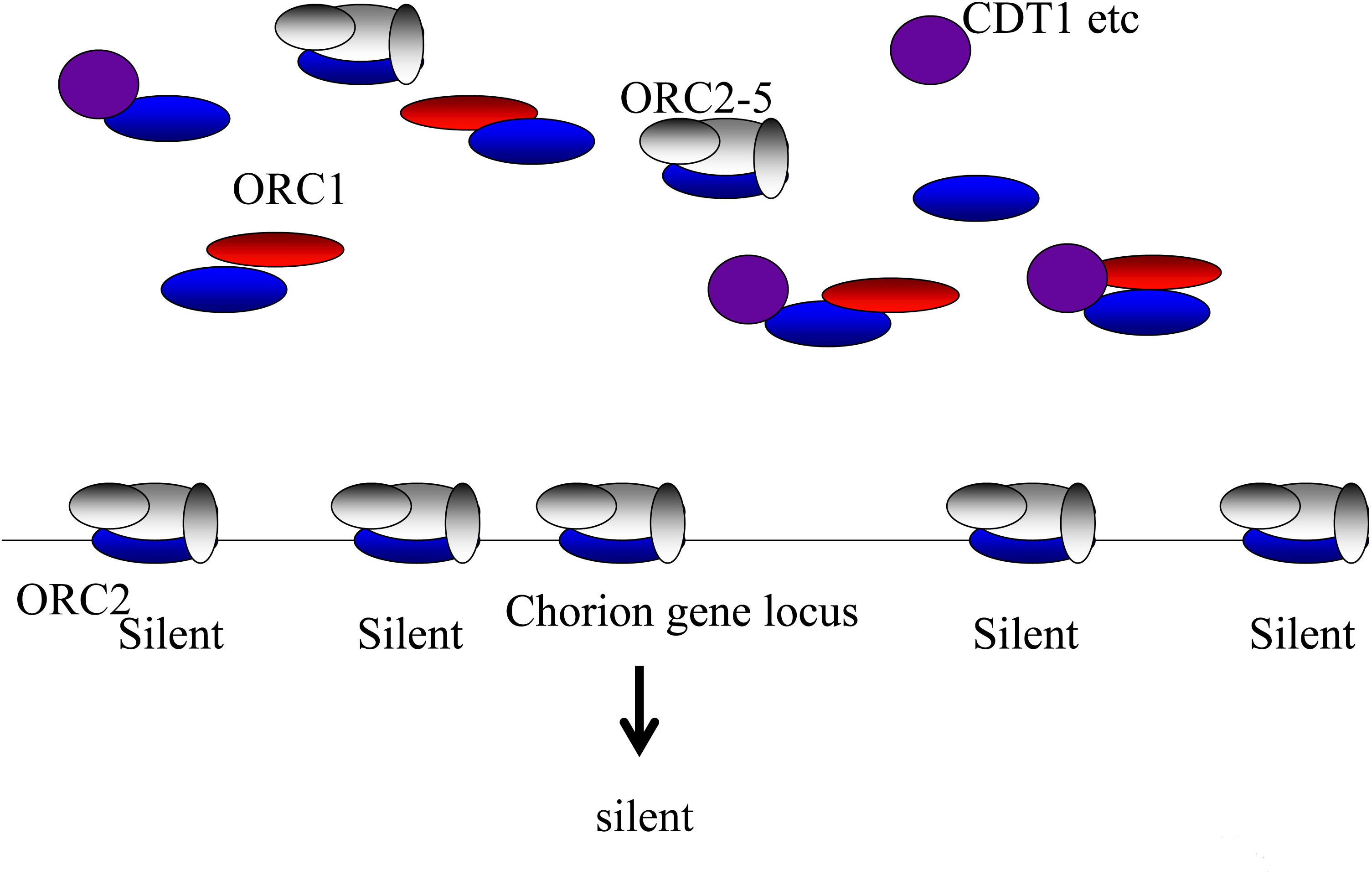
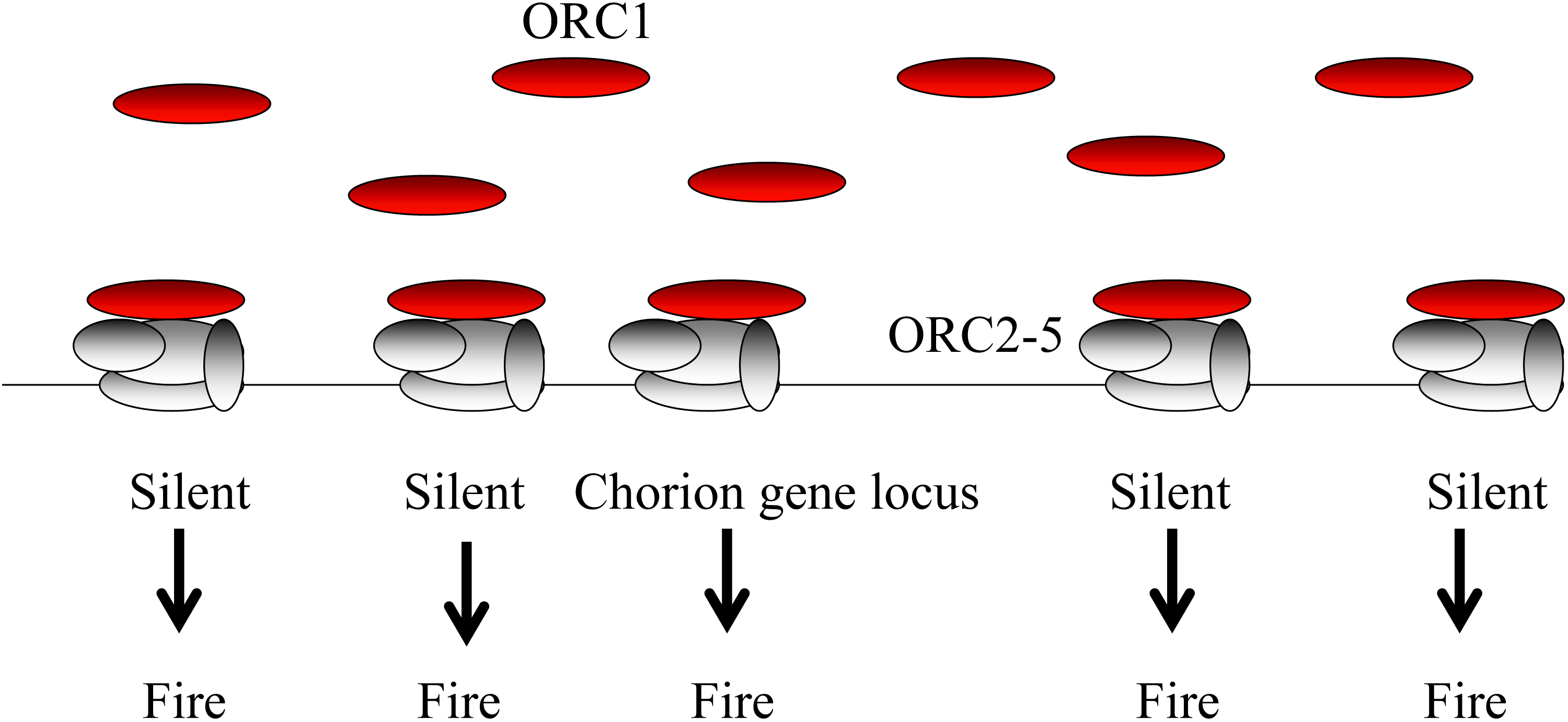
Schematic representation of the putative machanism of female sterile when Orc2 (A) and Orc1(B) was overexpressed in fc cells (Kohzaki et al., 2018a). (A) Overexpressed Orc2 titrate out the DNA replication machinery. (B) Overexpressed Orc1 disturbs programmed gene amplification.

The Ecdyson recepter (EcR) isoforms are functionally distinct. The early genes in tissues where the EcR-A isoform is dominant. Other EcR isoform, EcR-B1 is the predominant isoform in both the imaginal and larval cells of the larval midgut. What induce differences of expressions and functions? Bender et al. suggested that tissue-specific coactivators may provide the link between the transcription machinery for a given gene and a particular EcR isoform like TRR (Bender et al., 1997). In this situation, it would be the coactivator that determines which EcR isoform is used to activate the gene. These determinants might be akin to the plethora of putative coactivators recently found for vertebrate nuclear receptors (Mangelsdorf and Evans, 1995).

It is noteworthy that, in mammalian, many transcription factors turn out to be proto-oncogenes, including c-Jun, c-Myb and c-Myc. The oncogenecity of these oncogenes has been considered to arise from dysregulation of trasncription that they promote. We suspect, however, that dysregulation of replication caused by the maltifunction of these transcription factors contributes to their oncogenic potential. Several evidences have been increased (Murakami and Ito, 1999; Kohzaki and Murakami, 2005). In this issue, we showed that c-Jun could regulate the Orc loading in *S. cerevisiae* (Kohzaki at al., 1999). C-Jun orthologue, Gcn4 could recruite the Orc loading, too. A mutation in *Drosophila* myb gene includes a defect in progression of S phase in several tissues. Dominguez-Sola et al showed that c-Myc could modulate DNA replication origin activity through the regulation of Cdc45 loading (Dominguez-Sola et al., 2007; Lebofsky and Walter, 2007). We propose the DNA replication machinary contributes development (Kohzaki & Murakami, 2018b). These evidences may suggested that alterations of this space- and time-controlled process can lead to unscheduled DNA synthesis, checkpoint activation, genomic instability and /or cell death.

According to the Ministry of Health, Labor and Welfare of Japan, there are approximately 470,000 infertile individuals in Japan (Ministry of Health, Labor and Welfare of Japan HP). In America, the CDC (Center for Disease Control and Prevention, USA) reported the following related statistics:

- Percentage of women aged 15-44 with impaired fecundity: 12.1%
- Percentage of married women aged 15-44 that are infertile: 6.7%
- Number of women aged 15-44 who have ever used infertility services: 7.3 million
- Percentage of women aged 15-44 who have ever used infertility services: 12.0%

Infertility treatment, artificial insemination, and in vitro fertilization are routinely performed, and this is becoming a social problem. Infertility treatment, artificial insemination and in vitro fertilization are also performed, which is becoming a social problem. Screening strategies using these flies could also lead to the development of drugs that affect sterility and epigenetic effects related histone modification.

## Materials and Methods

### Fly strains

Fly strains were maintained at 25°C on standard food. *C323a-Gal4* driver and flis harboring UAS-EcR isoform transgene were obtained from the Bloomington Stock Center. Flis having *orc1*+-promoter-Orc1-GFP-9myc, UAS-Orc2-GFP-9myc, UAS-Orc1-GFP-9myc, UAS-Cdc6-GFP-9myc, UAS-GFP were from M. Asano (Duke University Medical Center). UAS-Lsd1 was a gift from N. Dyson (Harvard Medical School, Boston, USA). UAS-Lid and UAS-LidjmjC* were from R. N. Eisenman (Fred Hutchinson Cancer Research Center, Seattle, USA).

Femal sterile experiments were peformed as describe previously (Kohzaki et al., 2018a).

### ChIP assay

ChIP assay was performed mainly according to Austin et al (Austin et al., 1999; Kohzaki and Asano., 2016). Egg chambers from Orc1-GFP-9myc (Park and Asano, 1999) were dissected from ovaries of fattened flies in nonsupplimented Grace’s medium (GIBCO-BRL). Formaldehyde added to a final concentration of 2% and cross-linking was allowed to proceed for 15 min at room temperature on a rotator. The cross-linking reaction was stopped by addition of glycine at a final concentration of 0.125 mM and incubating 5 min. The cross-linked egg chamber were washed twice with 1 ml of TBS, then twice with 1ml of lysis buffer (Kohzaki and Murakami, 2007). The egg chambers were disrupted by sonication. Sonication and all postsonication procedures were performed as described previously (Kohzaki and Murakami, 2007). Myc antibody (9E10) was used in ChIP assay. EcR and TRR antibody was described previously (Sedkov et al, 2003). H3K4 Tri-Me (ab8580) and H3K4 Di-Me (ab7766) antibody are purchaced from abcom (England). Primers used in this issue were the same as those reported previously (Austin et al., 1999; Royzman et al., 1999).

### Microscope and Histolog

A Zeiss microscope and fluorescence was used to examine and photograph the ovaries.

Imaginal discs were fixed in either 1 or 2% paraformaldehyde (PFA) in PBS as previously described (Asano et al., 1996). Ovaries were fixed in 1% PFA in PBS and processed as described previously (Lin et al., 1994).

## Acknowledge

We thank N. Dyson (Harvard Medical School, Boston, USA), R. N. Eisenman (Fred Hutchinson Cancer Research Center, Seattle, USA) and M. Asano for flies harboring UAS-Lsd, UAS-Lid and UAS-LidjmjC*. We thanks M. Asano for technical advises and gifts of *orc1*+-promoter-Orc1-GFP-9myc, UAS-Orc2-GFP-9myc, UAS-Orc1-GFP-9myc, UAS-Cdc6-GFP-9myc, UAS-GFP. We dedicate this paper to M. Asano (The Ohio State University). We thank Dr. Tadashi Uemura (Kyoto University) and their lab for their dedicated support and helpful assistance. This work was partially supported by the Japanese Leukemia Research Fund. H.K. was supported by a KIT VL grant, the Memorial Fund on the 44 Meeting Annual of the Japan Society for Clinical Laboratory Automation and The Motoo Kimura Trust Foundation for the Promotion of Evolutionary Biology.

